# Ferritin-mediated iron homeostasis and bacterial shifts underpin drought adaptation in sorghum

**DOI:** 10.1101/2024.07.19.604343

**Authors:** Ahmad H. Kabir, Philip Brailey-Crane, Mostafa Abdelrahman, Jean Legeay, Bulbul Ahmed, Lam-Son Phan Tran, Jeffrey L. Bennetzen

## Abstract

Drought stress significantly impairs growth, and microbial interactions in sorghum. This study explores the transcriptional and microbial shifts in sorghum under drought, revealing key adaptations to water deficit. LC-MS (Liquid Chromatography-Mass Spectrometry) analyses revealed that drought stress induced abscisic acid while significantly reducing jasmonic acid levels in sorghum roots, likely due to resource conservation strategies during drought. Transcriptional reprogramming highlighted the upregulation of genes in the roots involved in mineral homeostasis (*Ferritin 1*, *Iron dehydrogenase*, *Nitrate transporter 1*), hormone signaling (*Ethylene-insensitive protein 3*, *Gibberellin 2-oxidase*), and osmotic regulation (*Aquaporin*, *Dehydrin*), underlining key adaptive responses to maintain nutrient uptake, redox status, and cellular turgor. In Fe-supplemented plants, increased Fe in roots correlated with increased *Ferritin 1* expression, improved plant health, and reduced Fenton reaction rate and H O levels. This suggests that ferritin helps minimize oxidative stress under drought in sorghum. Drought reduced root-associated bacterial diversity and richness while enriching drought tolerance-associated genera, such as *Burkholderia, Caballeronia* and *Paraburkholderia*, known for promoting plant growth through auxin production, phosphate solubilization, and siderophore-mediated iron acquisition. In contrast, fungal diversity and richness remained unchanged, dominated by *Talaromyces*, which showed a statistically non-significant increase under drought. Random forest models could not identify functional predictors for fungi but revealed a shift in bacterial functional groups under drought, with enrichment in phototrophy, methylotrophy, and nitrate reduction, traits emphasizing microbial roles in nutrient cycling and drought adaptation of sorghum. This study provides insights into the role of ferritin and potential bacterial bioinoculants that could enhance sorghum resilience to drought. Future research should validate these findings to integrate them into breeding programs and biofertilizer formulation for drought-tolerant sorghum and climate-resilient agriculture.

## Introduction

Drought stress, leading to adverse impacts on growth, development, and productivity (Lamaoui et al., 2018), is ubiquitous across crops and natural plant populations but varies dramatically in intensity and frequency. Primary drought stress symptoms, especially wilting and leaf curling, are reflective of reduced turgor pressure and water loss reduction strategies (Ruehr et al., 2019; Zoghi et al., 2019). Additional drought stress manifestations include leaf yellowing, stunted growth, and premature leaf senescence (Khan et al., 2018). Water deficiency disrupts essential physiological processes like photosynthesis, nutrient uptake, and hormone regulation, ultimately compromising the plant’s ability to thrive (Hussain et al., 2018). Additionally, crops subjected to drought stress often show increased susceptibility to pests and diseases (Duveiller et al., 2007). This major agricultural problem is expected to be more prevalent under current climate change scenarios. As global temperatures rise and weather patterns shift, many regions are experiencing unusually long periods of low precipitation and resultant water shortages. Sorghum (*Sorghum bicolor*) is known for its exceptional drought tolerance, but its yields are affected by prolonged periods of water scarcity (Rajarajan et al., 2021). Studies predict that drought, associated with temperature increases, could cause up to a 44% yield loss in US sorghum production in the near future, emphasizing the need for additional efforts to improve resilience, especially during the pre-flowering phase (Tack et al., 2017).

Plants employ diverse strategies to regulate water loss, including stomatal and osmotic adjustments through the accumulation of compatible solutes and the enhanced production of antioxidant enzymes and metabolites to counteract reactive oxygen species (ROS) generated under drought conditions (Seleiman et al., 2021; Takahashi et al., 2020). Hormones, particularly abscisic acid (ABA) and auxin, play a pivotal role in orchestrating these responses by modulating stomatal closure and activating stress-responsive genes, ultimately enhancing a plant’s ability to withstand periods of water shortage (Iqbal et al., 2022; Sadok and Schoppach, 2019). Minerals are also essential for plant drought tolerance, as they regulate osmotic adjustment, stomatal control, and stress signaling, which help plants maintain water balance (Khan et al., 2023). Particularly, calcium is vital for maintaining cell wall integrity and membrane stability, which can be compromised during drought conditions (Feng et al., 2023).

Microbes can improve plant stress resistance, including drought tolerance, by instigating a range of processes that alter plant biochemistry and physiology (Poudel et al., 2021; Timmusk et al., 2013). Fungal communities, including AMF (arbuscular mycorrhizal fungi), are sometimes associated with drought stress tolerance in some crops (Chen et al., 2021; Ferrol et al., 2019). Studies suggest that AMF can directly influence plant performance by modifying the plant–water relationship, leading to increased plant productivity during periods of drought stress (Ma et al., 2020; Rapparini and Peñuelas, 2014). *Trichoderma*, a fungal genus, has gained attention for its potential role in enhancing drought tolerance in plants. Two ways in which *Trichoderma* are believed to contribute to drought tolerance are by inducing systemic drought stress response in plants and by participating in the production of specific metabolites that positively impact plant responses to drought (Scudeletti et al., 2021; Mona et al., 2017). Monoderm bacterial lineages, such as *Actinobacteria*, *Chloroflexi*, and *Firmicutes*, induce increased desiccation tolerance in plants, associated with thicker plant cell walls, and these microbes’ increased abundances are driven by drought-responsive plant metabolism (Xu and Coleman-Derr, 2019). Inoculation of *Bacillus amyloliquefaciens* S-134 on wheat plants increased the production of auxins, stimulated root growth, and assisted plants in adapting to water stress conditions. Particularly, *Streptomyces* are reported to produce auxin and siderophores that can promote mineral availability, stress tolerance, and growth in plants (Anwar et al., 2016; Palaniyandi et al., 2014).

Sorghum is a promising bioenergy crop because of its high biomass yield and suitability for producing biofuels (Mathur et al., 2017; Prasad et al., 2021). Given the increasing frequency of drought events caused by climate change, understanding the molecular and microbial mechanisms underlying drought tolerance in sorghum is essential. Previous studies have documented individual mechanisms of drought tolerance, such as hormonal regulation or bacterial interactions (Qi et al., 2022; Rajarajan et al., 2021). However, the interconnected dynamics of plant transcriptomics and root-associated bacterial and fungal microbiomes remain underexplored, particularly in a bioenergy crop like sorghum at vegetative stage. Therefore, this research seeks to elucidate the interplay between plant gene expression and microbial interactions, identifying key gene networks and microbial taxa and their functional groups that may contribute to drought tolerance in sorghum.

## Materials and methods

### Plant cultivation and growth conditions

Seeds of sorghum (USDA-GRIN accession PI 591038) were surface sterilized using a 1% sodium hypochlorite bleach solution for 5 min. Following sterilization, the seeds were rinsed three times with sterile water and subsequently placed in a tray for germination at 25°C in an incubator for a duration of 2 d. Plants were cultivated in the greenhouse in pots in a mixture of field soil and SunGro potting mix (1:3) in two different conditions: control and drought. Drought conditions were initiated by reducing water availability to 75% compared to the controls (well-watered) two weeks after transplanting the seedlings into the soil pots. Daily monitoring of soil moisture content was conducted to maintain 75% and 25% soil moisture levels for control and drought-stressed plants, respectively. Plants were cultivated following a randomized complete block design (RCBD), with 9 replications (3 plants/pot) per treatment in a growth chamber (10 h light/14 h darkness, 250 μmol m ² s ¹) at 25°C for 6 weeks before measuring physiological properties and harvesting samples for other physiological and nucleic acid analyses, for all experiments. In an additional study, plants were cultivated for up to 4 weeks under four different treatments: control, drought, drought + FeEDTA (1 g/1000 g soil), and FeEDTA. Drought was imposed after two weeks.

### Determination of morphological and gas-exchange attributes

The height of each plant was measured using a measuring tape. The aboveground portions of the plant were harvested and dried for 3 days at 75°C in an incubator before obtaining the dry weight. Furthermore, the relative water content (%) in the leaf was measured as previously described (Soltys-Kalina et al., 2016). Relative water content (RWC) of leaves was measured on the fully expanded trifoliate leaves at the tip of the main stem as previously described (Schonfeld et al., 1988). Briefly, the fresh weights (FW) of leaves from each treatment were recorded immediately after removal. The turgid weight (TW) was measured after soaking the leaves in distilled water in beakers for 24 h at room temperature. Following the soaking period, the leaves were quickly and carefully blotted dry with tissue paper to prepare for turgid weight determination. The dry weight (DW) of the leaves was obtained by oven-drying the leaf samples for 72 h at 70°C. RWC was calculated using the formula: RWC (%) = (FW – DW) / (TW – DW) x 100.

Gas exchange and photosynthetic parameters of the leaves were measured using the LI-6800 portable Photosynthesis System (LI-COR Biosciences, USA). The photosynthetic rate, transpiration rate, stomatal conductance, and intrinsic water-use-efficiency (iWUE) were assessed on the central portion of the fourth fully expanded leaf under the following conditions: 2 cm² leaf area, flow rate of 600 μmol s ¹, relative humidity of 65%, CO_2_ reference of 400 μmol mol ¹, leaf vapor pressure deficit of 1.5 kPa, leaf temperature of 25°C, and leaf-level light intensity of 250 μmol photons m ² s ¹. Measurements were conducted three times, averaged for statistical analyses, and calculated as previously described (de Oliveira et al., 2022).

### Elemental analysis of plant tissues

After harvesting, the roots of 6-week-old sorghum plants were excised and rinsed with running tap water, followed by immersion in 0.1 mM CaSO_4_ for 10 min, and a final wash with deionized water. Subsequently, the roots and leaves were dried in an incubator for 3d at 75°C. Elemental concentrations were analyzed using ICP-MS (inductively coupled plasma mass spectrometry) at the Agricultural & Environmental Services Laboratories at the University of Georgia. The concentration of elements in the samples was determined by comparing the intensity of ion peaks in the spectrum to calibration standards.

### Determination of H_2_O_2_ and Fenton reaction rate in roots

For H_2_O_2_ determination, root samples were mixed in a solution of 0.1% trichloroacetic acid, as previously described (Alexieva et al., 2001). Briefly, the samples were finely ground into a powder using a mortar and pestle. The extracted fluid underwent centrifugation for 15 min at 10,000 rpm to separate cellular residues. The resulting upper aqueous phase was supplemented with potassium iodide (1 M) and phosphate buffer (10 mM, pH adjusted to 7.0), and then left in darkness for 60 min. Finally, the optical density of the solution was measured at 390 nm using a spectrophotometer (ThermoFisher, Waltham, USA), and the concentration of H_2_O_2_ was calculated based on the standard curve. Furthermore, the Fenton reaction rate in root samples was determined based on Fe and H O concentrations using the following formula (Wiegand et al. 2017): Fenton rate (µmol/s) = *k* × {Fe (µM)} × {H_2_O_2_ (µM), where *k* = 0.01 µM^-1^S^-1^ at pH 7 and 25°C.

### RNA-seq analysis in the roots

Before RNA extraction, roots were meticulously cleansed twice in sterile phosphate-buffered saline (PBS) through vortexing for 10 sec, followed by two rinses with sterile water to minimize surface contaminants. Subsequently, samples were pulverized into a powder using a pre-cooled pestle and mortar with liquid nitrogen. The resulting powdered material was then utilized for RNA extraction. Total RNA extraction was performed using the SV Total RNA Isolation System (Promega, USA), following the manufacturer’s instructions. A total of 1 µg of RNA per sample, possessing a RIN value >8, was employed as input material for RNA-seq library preparations. Sequencing libraries were constructed using the KAPA HiFi HotStart Library Amplification Kit (Kapa Biosystems, USA), per the manufacturer’s instructions. The resulting libraries were sequenced on the Illumina NextSeq SE 100bp platform. Overall, >80% of reads passed filters with Phred score values exceeding 33% per sample.

Adaptor sequences and low-quality sequences were respectively removed and filtered from raw reads using Trimmomatic v0.38 (Bolger et al., 2014). The sorghum reference genome (Sbicolor_454_v3.0.1) was obtained from Phytozome (McCormick et al., 2017) and the filtered reads were aligned to the reference genome using HISAT2 (v2.0.5) software. HT-Seq was utilized to count the read numbers mapped to each gene (Anders et al., 2014). Differentially expressed genes (DEGs) were identified using the Limma R package v3.19 (Law et al., 2016) and read counts were normalized by using the FPKM (fragments per kilobase of exon per million mapped fragments) method. Genes were identified as up-regulated or down-regulated using a cut-off threshold of log_2_ fold change (Log_2_FC) ≥ 1 and ≤ -1, respectively, with a false discovery rate (FDR) ≤ 0.05. Heatmaps were generated using pheatmap (Kolde, 2019) R package, respectively. Gene ontology for sorghum was retrieved using Phytozome (Goodstein et al., 2012) resources. Gene enrichment analysis was performed through the ShinyGO 0.80 web tool.

### Expression analysis of *Ferritin 1* in the roots

We performed RT-qPCR analysis to study the differential expression pattern of *Ferritin 1* in the roots of sorghum cultivated with or without FeEDTA and drought. B The first-strand cDNA was synthesized using the GoScript Reverse Transcription System (Promega, USA). Gene expression analysis was conducted via real-time PCR (qPCR) using the CFX96 Real-Time System (Bio-Rad, USA) using gene-specific primers (Fw: CCTCTTCGCCTACTTCGACC, Rv: AGAGCCAGCTCCATAGCGTA). The qPCR reaction conditions were 95°C for 3 min, 40 amplification cycles of 5 sec at 95°C, 30 sec annealing at 56°C, and an extension of 5 min at 60°C. Expression levels of target genes were determined using the dd^−ΔCt^ method (Livak and Schmittgen, 2001), where sorghum *GAPDH* (Fw: CCATCACTGCCACACAGAAAAC, Rv: AGGAACACGGAAGGACATACCAG) was used for normalization.

### Phytohormone analysis in roots

Phytohormone concentrations in sorghum roots were determined using LC-MS (Liquid Chromatography-Mass Spectrometry) at the Michigan State University Mass Spectrometry and Metabolomics Core. In brief, approximately 100 mg of fresh root material was homogenized in liquid nitrogen and extracted using a solution of 80:20 (v/v) methanol: water, 0.1% formic acid, and 0.1 g/L butylated hydroxytoluene. The samples were then placed in pre-frozen bead beater adaptors and beaten (30/s for 30 s) in a Tissue Lyser II (Qiagen, USA). Following a brief centrifugation (20 s) at 4°C to settle ground tissue at the tube bottom, the samples were further incubated on a shaker at 5,000 rpm at 4°C for 16 h. Subsequently, the samples were vortexed and centrifuged at 12,000 rpm for 10 min at 4°C, and the supernatant was transferred to autosampler vials. Standards for each phytohormone (Sigma-Aldrich, USA) were prepared in 80% methanol and 0.1% formic acid.

### Amplicon sequencing targeting fungi and bacteria

Harvested roots were cleaned by vortexing in sterile phosphate-buffered saline (PBS) for 10 seconds, followed by two rinses with sterile water to remove surface contaminants. Approximately 0.2 g aliquots of cleaned root samples were used for DNA extraction following the procedure outlined by Clarke (2009), along with RNase and Proteinase K treatments. Four biological replicates were used for microbiome analysis. Amplicon libraries were prepared for sequencing using a two-step PCR approach (Kozich et al., 2013). Two primer sets targeting regions of the ribosomal DNA gene were used to generate the libraries. The internal transcribed spacer (ITS) was used to target fungal communities (fITS7: GTGARTCATCGAATCTTTG and ITS4: TCCTCCGCTTATTGATATGC), whereas the small sub-unit encoding region (16S) was used to target bacterial communities (515F: TGTGYCAGCMGCCGCGGTAA and 806R: GGACTACNVGGGTWTCTAAT). Primers were adapted to include a 12N UDI sequence immediately prior to the target gene sequence to aid in cluster delineation when sequencing and Illumina P5/P7 complementary overhangs to facilitate barcode index addition in Step 2 PCR. Step 1 PCRs were carried out using Hot Start GoTaq master mixes (Promega, Madison, WI USA) in 12.5 ul volumes containing 2 ul of DNA, 2 mM dNTPs, and 10 pmol of each primer. PCR conditions were the same for both 16S and ITS amplicon generation: 95C for 5 min, 35 cycles at 94C for 30 s, 55C for 30s, 72C 30s, and 72C for 5 min. PCR steps were carried out using a BioRad T100 thermal cycler (Bio-Rad, Hercules, CA USA). Amplicons were dual barcoded with Illumina i7/i5 style barcodes in step 2 PCR using Phusion Hot Start II master mix (Thermo Fisher, Waltham, MA USA) to ensure high-fidelity barcode additions. Amplicons were sequenced using the NextSeq2000 platform on a P2 flow cell run in paired-end 300 bp mode (Illumina, San Diego, USA) by the Georgia Genomics and Bioinformatics Core (Athens, GA USA).

Adapter and primer sequences were trimmed from the raw sequencing reads using *Cutadapt v. 4.5* (Martin 2011). Next, low-quality reads were removed using the filterAndTrim function of *dada2* (*v. 1.28.0*) implemented in R (*v. 4.3.0*, Callahan et al., 2016), followed by error model estimation, dereplication, sample inference, and nonchimeric removal to generate a table of amplicon sequence variants (ASVs). Prior to taxonomy assignment, ASVs were pre-filtered by BLAST-matching against the UNITE database for ITS (dynamic_s_all_25.07.2023, Abarenkov et al., 2024) and the SILVA database for 16S (v 138.2_SSURef_NR99, Quast et al., 2013). Any sequences with a query coverage < 80% were removed. Both ITS and 16S taxonomy was assigned using CONSTAX2 (Liber *et al.,* 2021), which carries out consensus taxonomy assignment from multiple approaches, using the same databases. Further taxonomic curation was performed using *phyloseq* implemented in R (Callahan et al., 2016). 16S OTUs were filtered to include only OTUs assigned to the Bacterial kingdom as the ‘higher level taxonomy’ assignment, which excludes chloroplast and mitochondrial sequences. ITS OTUs were filtered to remove any OTUs where the ‘higher level taxonomy’ assignment through CONSTAX2 was not to the Fungal kingdom. Alpha diversity metrics (observed species richness and Shannon index) were calculated using *phyloseq*. Alpha diversity metrics were compared between treatment groups using ANOVA, followed by post-hoc testing where appropriate with ‘hsd’ multiple testing corrections employed. Beta diversity was assessed through Bray-Curtis dissimilarities calculated between samples and statistically compared between treatments with PERMANOVA analysis using vegan (*v. 2.6.4*). Community dissimilarity between samples and treatments was visualized through principal co-ordinate analysis (PCoA). Two methods were employed to determine which taxa were more associated with different treatments at the genus level. Indicator species analysis was conducted using the R package indicspecies (*v. 1.7.14*) on a core set of genera (>0.001 RA in > 10% of samples) consisting of 80 out of 306 genera. The resulting *p*-values were adjusted for multiple testing using “fdr” correction. The relative abundances of the top ten most abundant genera of bacteria and fungi were also compared between treatments using ANOVA, again followed by post-hoc testing where appropriate.

Furthermore, the dataset was trimmed to retain only taxa accounting for more than 0.1% of the total reads before conducting co-occurrence network analyses and random forest modeling. Co-occurrence networks were calculated and plotted using SpiecEasi (Kurtz et al., 2015) and igraph (Antonov et al., 2024) packages in R, with betweenness centrality values computed for all ASVs using the betweenness function of igraph. Taxa in the top 5% of betweenness values were designated as hub taxa. Random forest models, implemented using the randomForest (Breiman 2001) package in R with 100 iterations of decision trees, were used to identify ASVs predictive of drought conditions, with ASVs ranked by their GINI index, where a lower index indicates higher predictive importance. Functional pathways for bacterial 16S rRNA ASVs were attributed using the PICRUSt2 (Langille et al., 2013) package and the MetaCyc (Karp et al., 2002) database, while bacterial functional traits were assigned using the FAPROTAX (Louca et al., 2016) package. Similarly, fungal ITS ASVs were annotated with functional groups using the FunFun (Krivonos et al., 2023) package and the KEGG (Kanehisa et al., 2000) database.

### Statistical analysis of plant characteristics

All measurements were conducted with at least three independent biological replicates for each treatment. To identify significant differences between treatment groups (*P* < 0.05), student t-test and analysis of variance (ANOVA) followed by Duncan’s Multiple Range Test (DMRT) were carried out with treatment as an independent variable using SPSS Statistics 20.0 (IBM). Graphical representation was performed using GraphPad Prism 6.0.

## Results

### Changes in morphological and physiological traits under drought stress

Drought stress caused a substantial decline in shoot and root systems compared to controls in these growth chamber experiments, with soils containing natural field microbiomes (Fig. 1A and 1B). The root length, root dry weight, shoot height, shoot dry weight, leaf relative water content, and leaf chlorophyll score showed a significant decrease under drought compared to controls (Fig. 1C-1H). Furthermore, we found a significant increase in H_2_O_2_ concentrations in the roots of sorghum exposed to drought relative to controls (Fig. 1I).

**Fig. 1.**
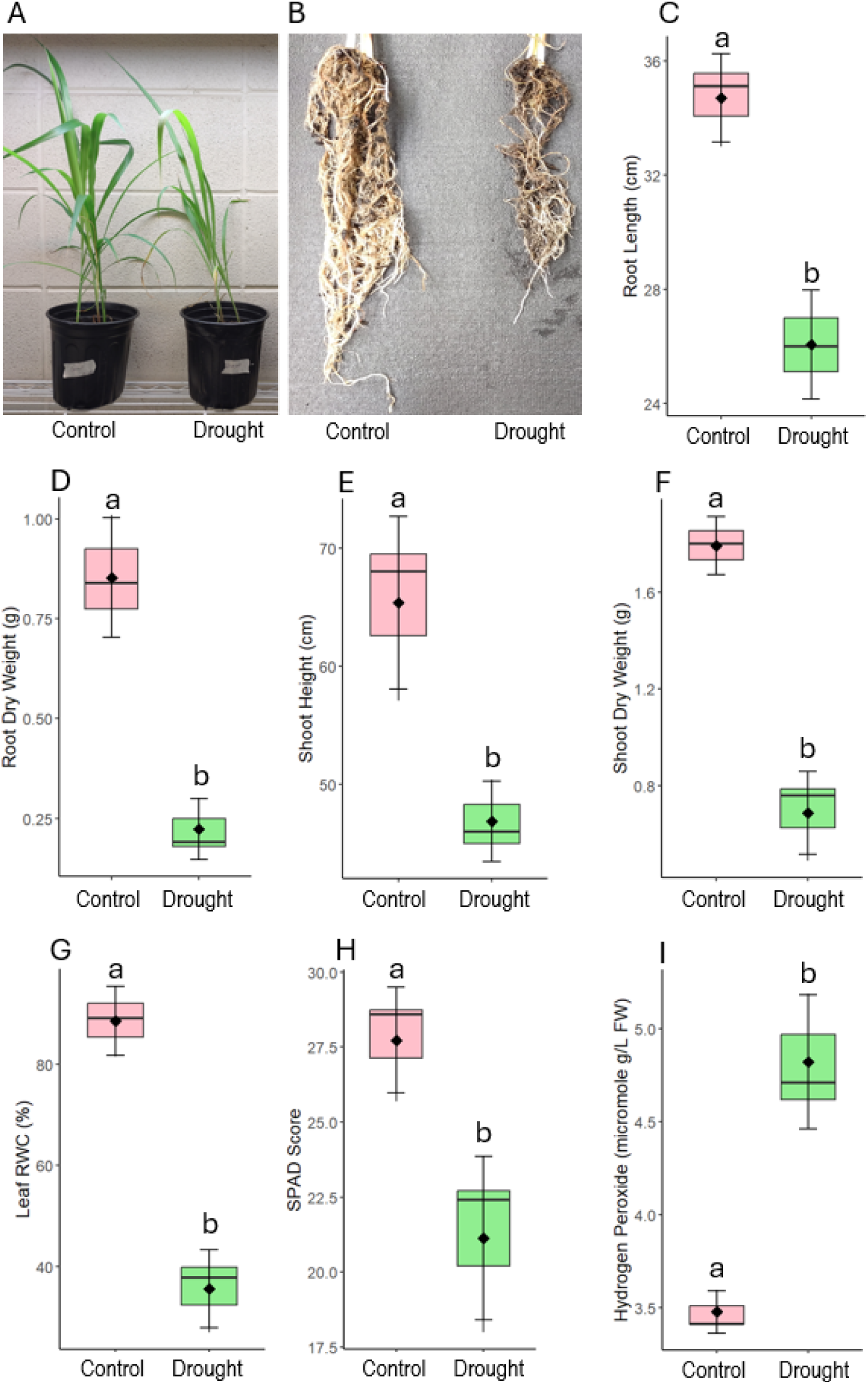
Changes in morphological and physiological attributes in sorghum cultivated with or without drought and Fe supplementation. Shoot phenotype (A), root phenotype (B), root length (C), root dry weight (D), shoot height (E), shoot dry weight (F), leaf relative water content (G), leaf SPAD score (H) and root H_2_O_2_ concentration (I) in sorghum cultivated in the absence (control) or presence of drought. The data presented are means with standard deviations (*n* = 3) of individual replicates). Statistically significant differences among the treatments at a *p* < 0.05 level are indicated by different alphabetical letters (a, b) above the bars, where applicable. Plants were cultivated in a growth chamber for 6 weeks.

ICP-MS was employed to investigate changes in elemental concentrations in the roots and shoots of sorghum plants. Fe concentration was significantly different between treatments, exhibiting a significant decline in the roots and shoots of drought-exposed sorghum (Table 1). Zn level significantly reduced in both root and shoot with drought. Mn levels significantly declined in the roots but showed no significant alternations in the shoot with drought compared to controls. S levels significantly declined in both root and shoot with drought compared to controls. Ca concentration significantly increased in the roots of drought-imposed plants relative to controls (Table 1).

**Table 1.** ICP-MS analysis of nutrients in the root and shoot of sorghum grown in control and drought conditions. Different letters indicate significant differences between the control and high pH conditions at a *P* <0.05 level, based on *t*-test. The data represents the means ± SD of three independent biological samples (*n* = 3). Plants were cultivated in a growth chamber for 6 weeks.

| Elements | Root |  | Shoot |  |
| --- | --- | --- | --- | --- |
|  | Control | Drought | Control | Drought |
| Fe (ppm) | 395±75<sup>a</sup> | 305±61<sup>b</sup> | 156±31<sup>a</sup> | 96±7.76<sup>b</sup> |
| Zn (ppm) | 47±5.6<sup>a</sup> | 40±4.5<sup>a</sup> | 62±9.9<sup>a</sup> | 59±17.1<sup>a</sup> |
| Mn (ppm) | 52±19<sup>a</sup> | 24.1±7.2<sup>b</sup> | 40±2.8<sup>a</sup> | 47.1±11.9<sup>a</sup> |
| S (%) | 0.05±0.015<sup>a</sup> | 0.025±0.005<sup>b</sup> | 0.034±0.007<sup>a</sup> | 0.024±0.003<sup>b</sup> |
| Ca (%) | 0.16±0.04<sup>a</sup> | 0.33±0.06<sup>b</sup> | 0.82±0.34<sup>a</sup> | 0.92±0.05<sup>a</sup> |

### Changes in leaf gas-exchange parameters and root phytohormone

We measured the gas-exchange parameters in the leaves of sorghum cultivated under different treatments. In this study, drought caused a significant decrease in the photosynthetic rate, transpiration rate, stomatal conductance, and water use efficiency (intrinsic) compared to well-watered controls (Table 2). Furthermore, drought-exposed plants showed a significant decrease in IAA concentrations in roots relative to controls (Table 2). Sorghum plants exposed to drought showed a significant increase in ABA concentrations in roots in contrast to control plants with added Fe boosting the ABA level even higher. Relative to well-watered controls, drought caused a significant decline in JA concentrations in the roots of sorghum (Table 2). In this study, GA3 levels showed no significant changes in the roots of sorghum cultivated under different treatments of drought and Fe supplementation (Table 2).

**Table 2.** Changes in leaf gas-exchange parameters and root phytohormone in sorghum cultivated in control and drought conditions. Different letters in superscript within a row (i.e. phenotype) indicate significant differences between means ± SD of three independent biological replicates (*n* = 3) at *p* < 0.05 based on *t*-test. Plants were cultivated in a growth chamber for 6 weeks.

| Parameters | Control | Drought |
| --- | --- | --- |
| Photosynthetic rate ($\mu\text{mol m}^{-2} \text{s}^{-1}$) | 19±1.3<sup>a</sup> | 5±0.52<sup>b</sup> |
| Transpiration rate ($\mu\text{mol m}^{-2} \text{s}^{-1}$) | 0.002±0.0009<sup>a</sup> | 0.0005±0.0001<sup>b</sup> |
| Stomatal conductance ($\mu\text{mol m}^{-2} \text{s}^{-1}$) | 0.09±0.01<sup>a</sup> | 0.005±0.003<sup>b</sup> |
| Water use efficiency (intrinsic) ($\mu\text{mol mol}^{-1}$) | 195±12.8<sup>a</sup> | 72±2.9<sup>b</sup> |
| Plant hormone | Control | Drought |
| IAA ($\text{ng g}^{-1}$) | 8.3±1.5<sup>a</sup> | 4.4±1.0<sup>b</sup> |
| ABA ($\text{ng g}^{-1}$) | 3.7±1.4<sup>a</sup> | 78.6±28.9<sup>b</sup> |
| JA ($\text{ng g}^{-1}$) | 34.5±1.4<sup>a</sup> | 9.2±2.5<sup>b</sup> |
| GA3 ($\text{ng g}^{-1}$) | 10.5±2.4<sup>a</sup> | 9.3±2.2<sup>a</sup> |

### RNA-seq analysis of differentially expressed genes in roots of sorghum

Transcript abundances in sorghum roots were seen to be quite different for the two treatments (Fig. 2). Principal component analysis (PCA) based on three biological replicates revealed that the first two principal components, PC1 and PC2, contributed 51.5% and 19.2% of the total variance, respectively (Fig. 2A). Control plants clustered far from the drought-exposed samples. The analysis of differentially expressed genes (DEGs) in sorghum roots showed significantly different gene activity in response to two different conditions (Fig. 2B). There were 640 upregulated (2B) and 744 downregulated (2C) genes in ‘drought vs. control’ comparison (Fig. 2B).

**Fig. 2.**
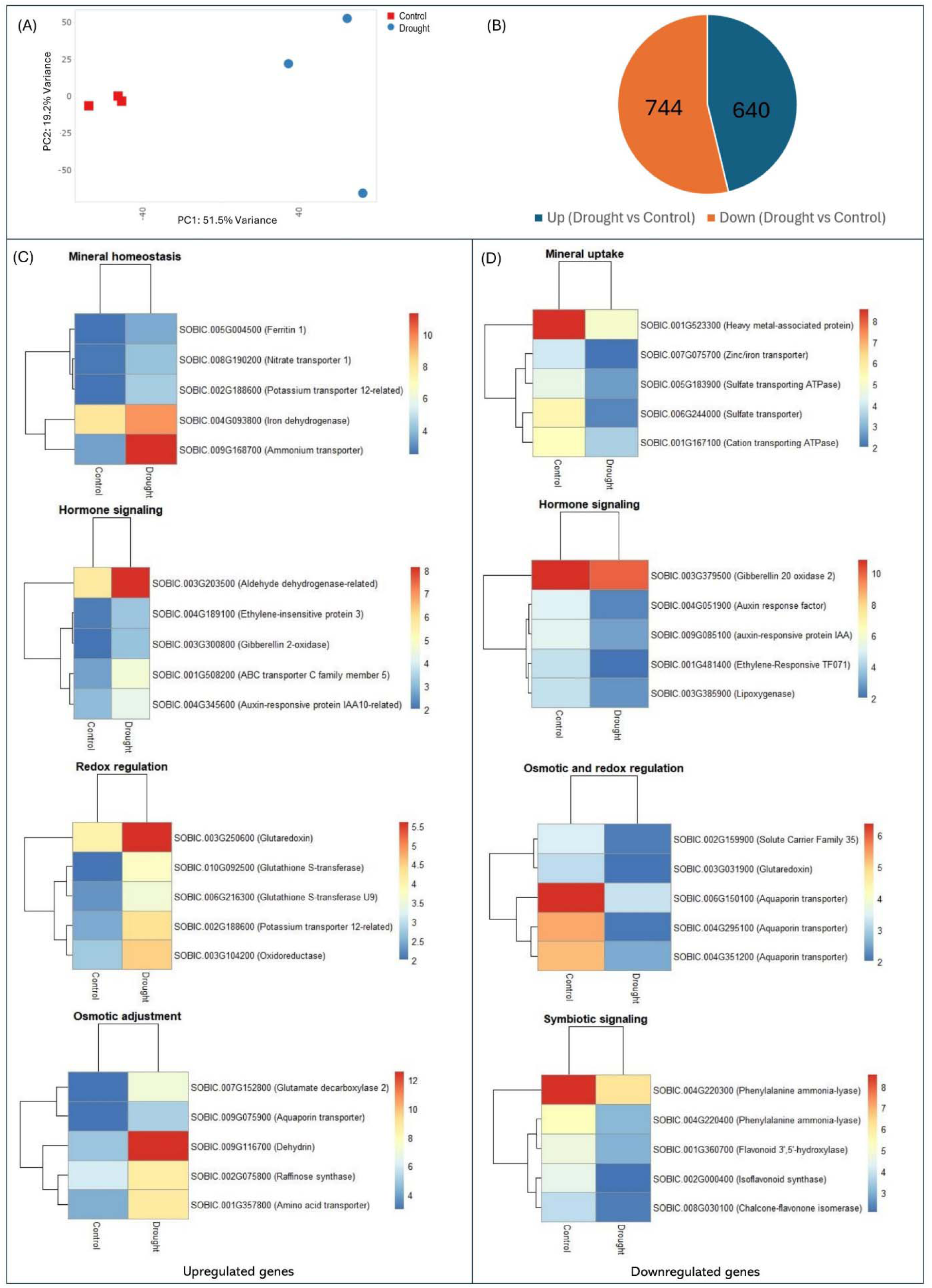
Principal component analysis of samples (A), Pie chart of differentially expressed genes (> Log2 fold, p value < 0.05) (B), top upregulated genes (C) and top downregulated genes (D) in the roots of sorghum plants cultivated in controla and drought conditions. A minimum of 20 million reads on average, with more than 33% read quality per sample, were utilized per treatment. Plants were cultivated in a growth chamber for 6 weeks.

We further annotated the functions of DEGs between control and drought using the Phytozome (V. 13) database for sorghum. We have categorized several differentially expressed (*P* < 0.05) genes known to be associated with drought and stress responses in earlier plant studies for more detailed investigation. Drought imposed to plants caused a significant upregulation of several crucial genes related to mineral homeostasis (*Sobic.005G004500 - Ferritin 1*; *Sobic.008G190200 - Nitrate transporter 1*; *Sobic.002G188600 - Potassium transporter 12-related*; *Sobic.004G093800 - Iron dehydrogenase*; *Sobic.009G168700 - Ammonium transporter*), osmotic adjustment (*Sobic.007G152800 - Glutamate decarboxylase 2*; *Sobic.009G075900 - Aquaporin transporter*; *Sobic.009G116700 - Dehydrin*; *Sobic.002G075800 - Raffinose synthase*; *Sobic.001G357800 - Amino acid transporter*), hormone signaling (*Sobic.003G203500 - Aldehyde dehydrogenase-related*; *Sobic.004G189100 - Ethylene-insensitive protein 3*; *Sobic.003G300800 - Gibberellin 2-oxidase*; *Sobic.001G508200 - ABC transporter C family member 5*; *Sobic.004G345600 - Auxin-responsive protein IAA10-related*) and *redox regulation (Sobic.003G250600 - Glutaredoxin*; *Sobic.010G092500 - Glutathione S-transferase*; *Sobic.006G216300 - Glutathione S-transferase U9*; *Sobic.002G188600 - Potassium transporter 12-related*; *Sobic.003G104200 - Oxidoreductase*) in the roots (Fig. 2C). In addition, several genes associated with mineral uptake (*Sobic.001G523300 - Heavy metal-associated protein*; *Sobic.007G075700 - Zinc/iron transporter*; *Sobic.005G183900 - Sulfate transporting ATPase*; *Sobic.006G244000 - Sulfate transporter*; *Sobic.001G167100 - Cation transporting ATPase*), hormone signaling (*Sobic.003G379500 - Gibberellin 20 oxidase 2*; *Sobic.004G051900 - Auxin response factor*; *Sobic.009G085100 - Auxin-responsive protein IAA*; *Sobic.001G481400 - Ethylene-responsive TF0171*; *Sobic.003G385900 - Lipooxygenase*), osmotic and redox regulation (*Sobic.002G159900 - Solute carried family 35*; *Sobic.003G031900 - Glutaredoxin*; *Sobic.006G150100 - Aquaporin transporter*; *Sobic.004G295100 - Aquaporin transporter*; *Sobic.004G351200 - Aquaporin transporter*), and symbiotic signaling (*Sobic.004G220300 - Phenylalanine ammonia-lyase*; *Sobic.004G220400 - Phenylalanine ammonia-lyase*; *Sobic.001G360700 - Flavonoid 3’,5’-hydroxylase*; *Sobic.002G000400 - Isoflavonoid synthase*; *Sobic.008G030100 - Chalcone-flavonone isomerase*) were found to be downregulated in response to drought (Fig. 2D).

We have analyzed the biological significance of DEGs in roots of sorghum in ‘control vs drought’ conditions. Several pathways with genes that were highly enriched in representation within the upregulated DEGs included, but were not limited to, those involved in various biological processes (response to hydrogen peroxide, protein complex oligomerization, response to heat, response to ROS, response to osmotic stress, response to abscisic acid, response to abiotic stimulus, response to oxygen-containing compound) and KEGG pathways (galactose metabolism, carotenoid biosynthesis, MAPK signaling pathway, amino sugar and nucleotide sugar metabolism, metabolic pathways) in sorghum (Fig. 3A). Conversely, downregulated genes were enriched in pathways related to biological processes such as nucleosome assembly, chromatin organization, cellular protein-containing complex assembly, peptide biosynthetic process, cellular component biogenesis, organic substance biosynthetic process) and various KEGG pathways (cyanoamino acid metabolism, ribosome, phenylalanine metabolism, glutathione metabolism, phenylpropanoid biosynthesis, biosynthesis of amino acid, biosynthesis of secondary metabolites) (Fig. 3B).

**Fig. 3.**
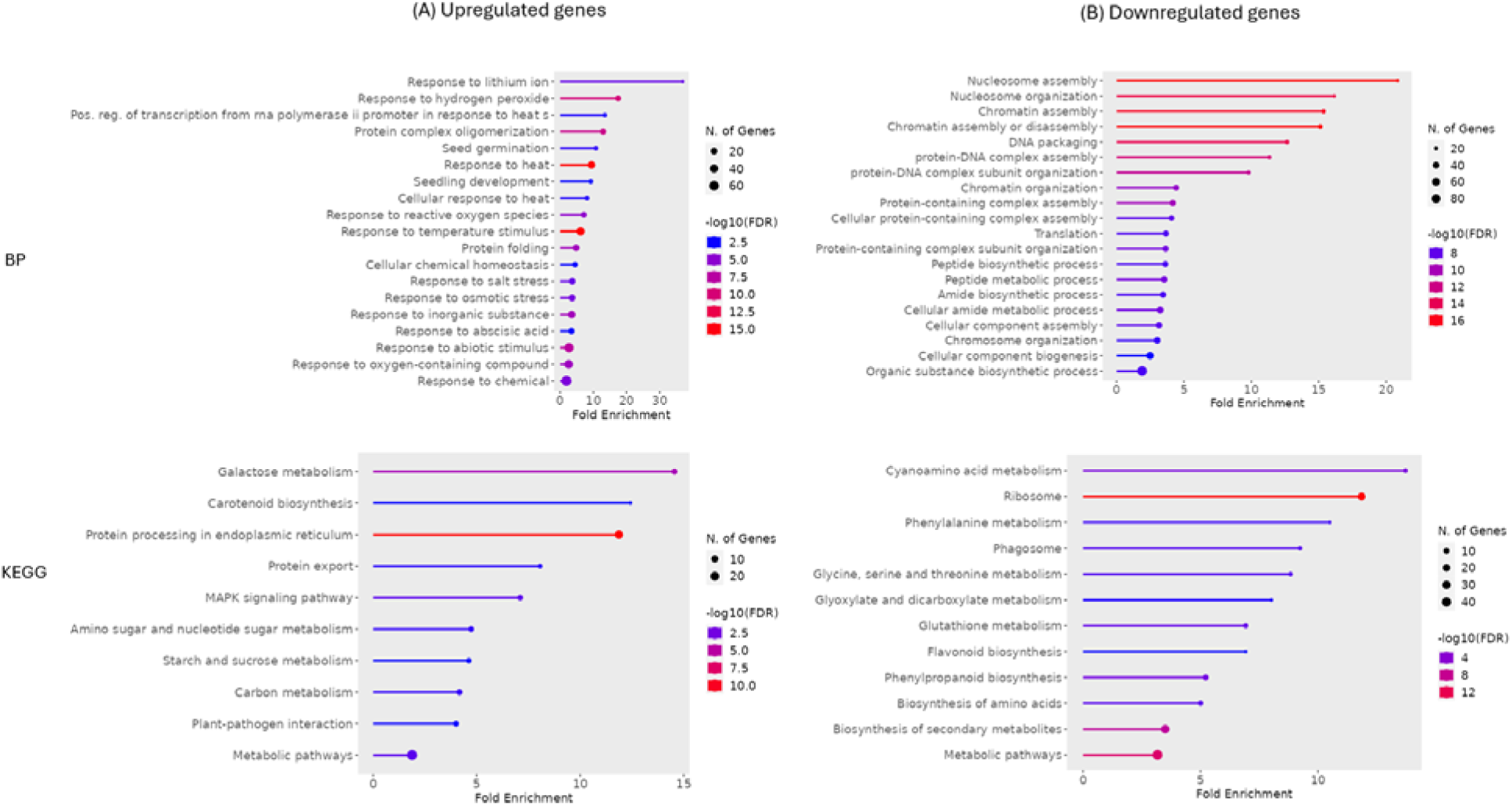
Gene ontology and enrichment analysis of pathways classified as biological process (BP), and KEGG pathways. These pathways or terms within the list of genes differentially (>Log2 fold, p value <0.05) upregulated (A) and downregulated (B) in ‘drought vs control’ conditions in roots of sorghum.

### Changes in diversity and composition of bacterial and fungal communities

Total bacterial and fungal communities in sorghum roots were analyzed by ribosomal DNA amplicon sequencing. PERMANOVA analysis of bacterial community composition revealed that drought significantly altered the composition of the bacterial (F = 6.3682, df = 1, p = 0.03) and fungal (F = 7.7593, df = 1, p = 0.03) communities (Fig. 4A). The experimental drought treatment was found to significantly alter the observed OTU richness for bacteria (*F* = 72.28, *df* = 1, *p* = 0.000145) but not for fungi (*F* = 3.667, *df* = 1, *p* = 0.104) in the roots (Fig 4B). Similarly, the Shannon evenness of bacterial (*F* = 324, *df* = 1, *p* = 1.89e-06) communities showed a significant change, while fungi (*F* = 1.829, *df* = 1, *p* = 0.225) remained unchanged (Fig. 4B).

**Fig. 4.**
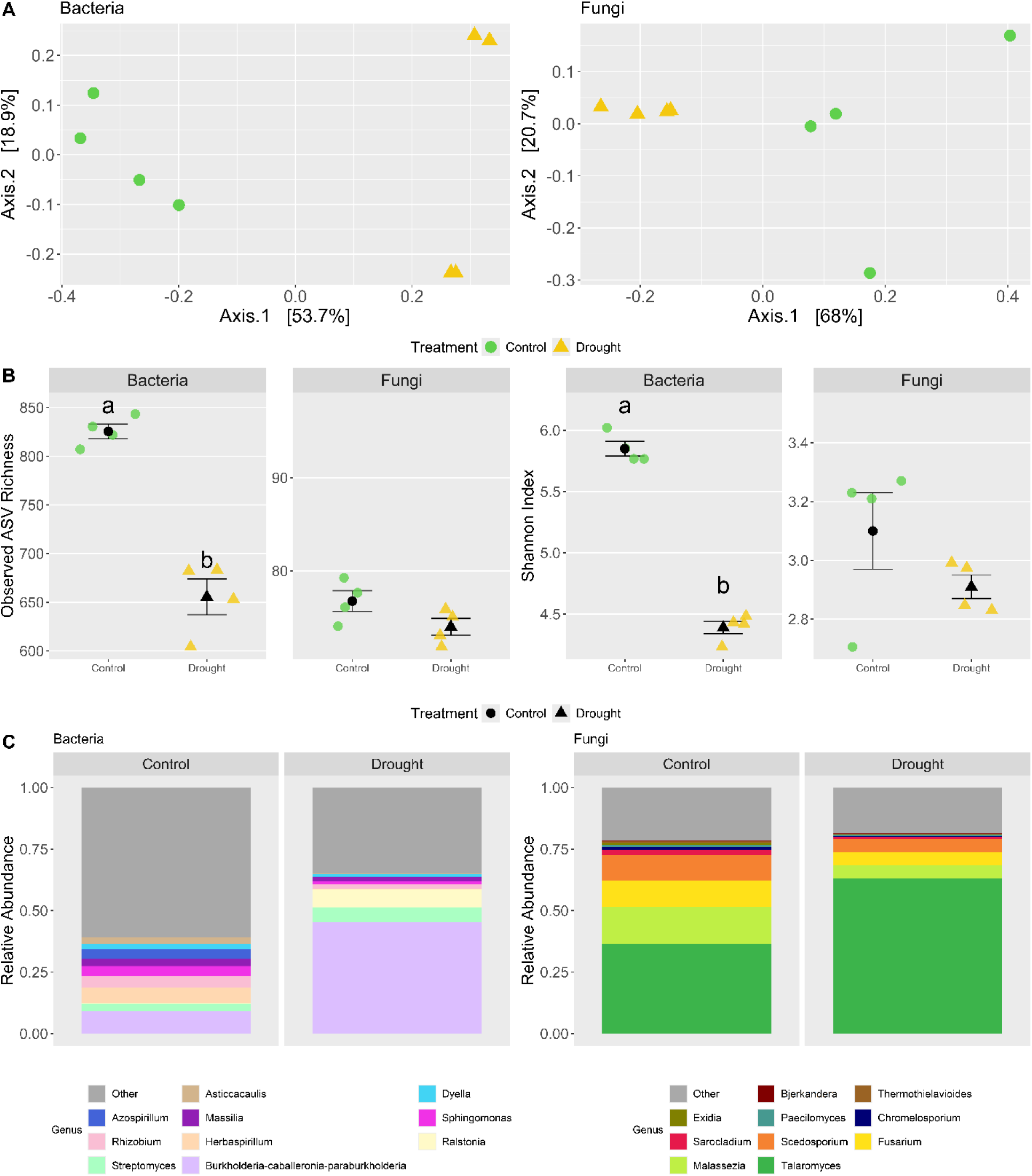
Panel A) Non-metric dimensional scaling (NMDS) ordination that displays community dissimilarity between samples for bacterial and fungal communities among the treatments. Mean centroids are displayed as diamonds ± standard errors in the NMDS1 and NMDS2 axes to demonstrate the dispersal of data points around the mean within each treatment group. Individual sample shapes and colors are associated with experimental treatment (green / circle = control, yellow / triangle = drought). Panel B) Observed OTU richness and Shannon index values for bacterial and fungal communities. Mean values ± SE for each treatment is displayed in black, and individual sample values are colored by treatment. Letters denote distinct statistical groupings of treatments determined through post-hoc pairwise testing. Panel C) Stacked barplots displaying the mean relative abundance of the top 10 most abundant genera for bacteria and fungi. The ‘Other’ category includes both lower abundance genera and taxa that were unclassified at the genus level.

Differential abundance tests were performed at the genus level for the top ten most abundant genera to balance the specificity of taxa interactions, statistical power of the analysis, and redundancy, as the majority of OTUs in both datasets are single representatives of their genera. Four of the ten most abundant bacterial genera were found to be differentially abundant between treatments. In particular, *Burkholderia-Caballeronia-Paraburkholderia* was in very low abundance in the control samples and increased through drought. However, the relative abundance of *Herbaspirillum*, *Rhizobium,* and *Sphingomonas* were negatively affected by the drought treatment, exhibiting lower relative abundances than control samples (Fig. 4C). We found that *Azospirillum*, *Streptomyces*, *Asticcacaulis*, *Massilia*, *Dyella,* and *Ralstonia* remained unchanged due to drought. In the case of fungi, only *Malassezia* showed a significant increase while the other nine (*Exidia*, *Sarocladium*, *Bjerkandera*, *Paecilomyces*, *Scedosporium*, *Talaromyces*, *Thermothielavioides*, *Chromelosporium* and *Fusarium*) among the top ten representative genera showed no significant shift in the roots due to drought (Fig. 4C).

There was no effect of drought on the general functional composition of either bacteria (*p* = 0.058) or fungi (p = 0.942). Random forest models were not able to find functional predictors for fungi (an error rate of 88%) but were able to find functional predictors for bacteria (an error rate of 12.5%). We also identified the functional groups associated with bacterial communities under control or drought conditions, predicted using either the MetaCyc (M) database or the FAPROTAX (F) annotation (Table 3). Most groups, such as chorismate metabolism, glycolysis, L-lysine biosynthesis II, and protochatechuate degradation I, were associated with the control condition. In contrast, groups like clavulanate biosynthesis, phototrophy, and photosynthetic cyanobacteria were specifically linked to drought conditions. While the majority of functional predictions were derived from the MetaCyc database, FAPROTAX contributed to identifying additional drought-associated groups, including methyltrophy and nitrate reduction. This highlights the differential functional responses of bacterial communities under varying environmental conditions (Table 3).

**Table 3.** The best functional predictors associated with either drought or control were assigned using the MetaCyc pathway database or the FAPROTAX functional annotation.

| Metacyc label (M) or Faptraox prediction (F) | Pathway | Treatment group |
| --- | --- | --- |
| ALL-CHORISMATE-PWY(M) | Chorismate metabolism | Control |
| GLYCOLYSIS(M) | Glycolysis | Control |
| P184-PWY(M) | Protocatechuate degradation I | Control |
| P341-PWY(M) | Pyrococcus | Control |
| PWY-2941(M) | L-lysine biosynthesis II | Control |
| PWY-5121(M) | Geranylgeranyl diphosphate biosynthesis II | Control |
| PWY-5896(M) | Menaquinol-10 biosynthesis | Control |
| PWY-5679(M) | Clavulanate biosynthesis | Drought |
| PWY-6071(M) | Phenylethylamine degradation | Control |
| (F) | Nitrate reduction | Control |
| (F) | Methylotrophy | Control |
| (F) | Phototrophy | Drought |
| (F) | Photosynthetic Cyanobacteria | Drought |

### Effect of Fe supplement in drought tolerance

The visual comparison of sorghum plants under different treatments showed noticeable differences in growth. Control and FeEDTA-treated drought-stressed plants appeared healthier and more vigorous than drought-stressed plants without FeEDTA (Fig. 5A). Drought significantly reduced chlorophyll content (SPAD values), but FeEDTA treatment increased SPAD scores in both control and drought-exposed plants (Fig. 5B). Leaf relative water content (RWC) was lower in drought-stressed plants compared to controls. However, FeEDTA application under drought conditions showed a significant increase in leaf RWC compared to control plants (Fig. 5C). Drought-stressed plants showed significantly reduced shoot height compared to control plants (Fig. 5D). However, plants treated with FeEDTA under drought conditions exhibited the highest shoot height, significantly greater than both control and drought-exposed plants. FeEDTA applied to drought conditions showed a significant increase in shoot height compared to the plants solely cultivated under drought (Fig. 5D). Also, drought-stressed plants showed significantly reduced shoot dry weight compared to control plants (Fig. 5E). However, plants treated with FeEDTA under drought exhibited a significant increase in shoot dry weight compared to drought-exposed plants. FeEDTA applied under control conditions showed the highest shoot dry weight compared to other treatment groups (Fig. 5E). Furthermore, the root length and root dry weight showed no significant changes in sorghum with or without drought in the absence or presence of FeEDTA compared to controls plants (Fig. 5F-5G).

**Fig. 5.**
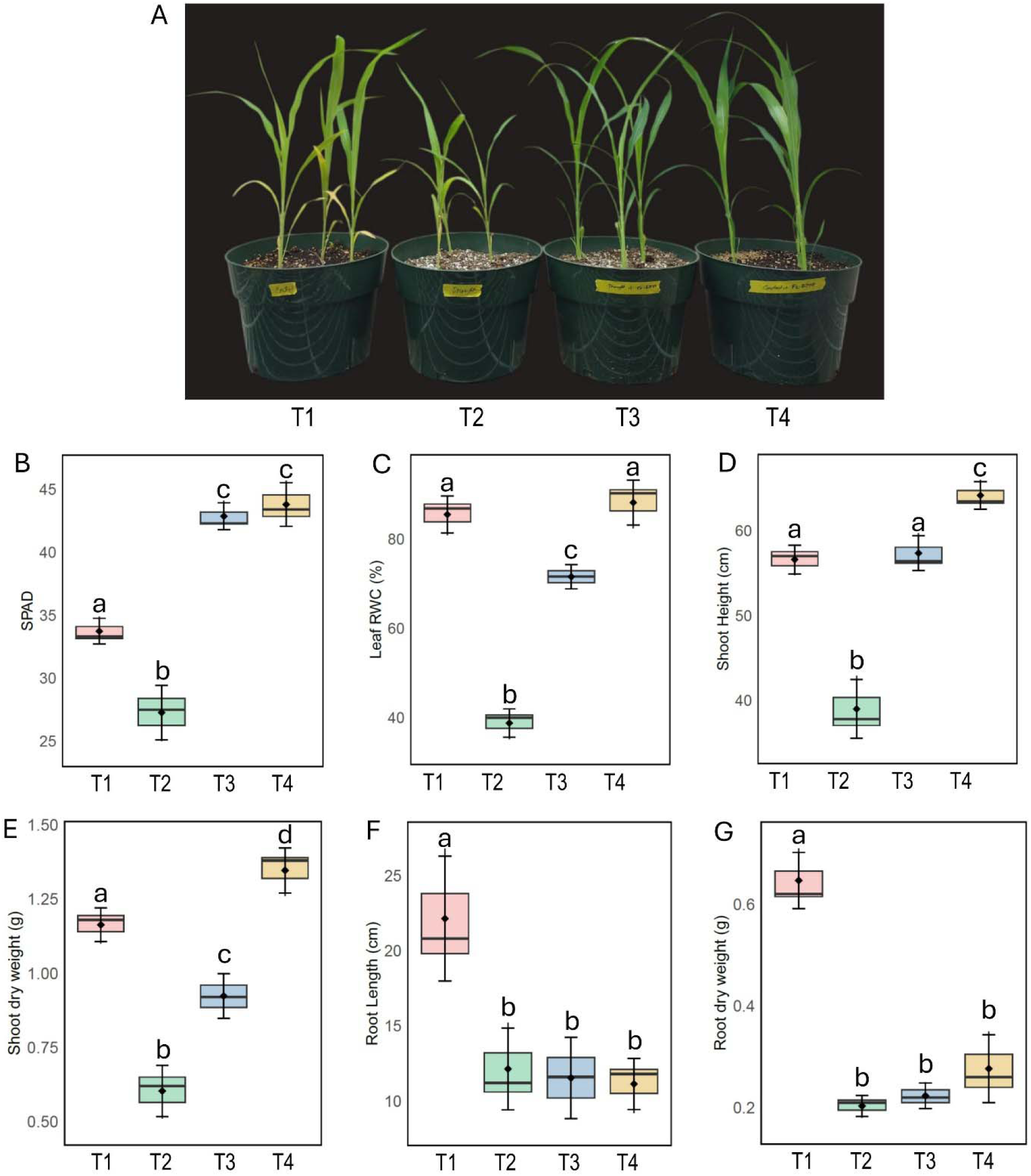
Changes in aboveground phenotype (A), SPAD score (B), leaf relative water content (C), shoot height (D), shoot dry weight (E), root length (F) and root dry weight (G) in sorghum cultivated with different drought and Fe treatments (T1: control, T2: drought, T3: drought + FeEDTA, T4: FeEDTA). The data presented are means with standard deviations (*n* = 3) of individual replicates). Statistically significant differences among the treatments at a *p* < 0.05 level are indicated by different alphabetical letters above the bars, where applicable. Plants were cultivated in a growth chamber for 4 weeks.

Drought stress significantly reduced root and shoot Fe levels compared to control plants. However, FeEDTA application restored Fe levels under both control and drought conditions (Fig. 6A-6B). Oxidative stress markers were also affected, with drought-stressed plants showing higher H_2_O_2_ levels, while FeEDTA application under drought conditions significantly reduced H_2_O_2_ in the roots (Fig. 6C). Fenton reaction rate showed a significant increase in the roots of sorghum under drought compared to controls. Interestingly, FeEDTA added to drought-stressed plants showed a significant decline in Fenton reaction compared to drought-exposed plants. FeEDTA supplemented with control plants showed a significantly higher Fenton reaction rate compared to controls (Fig. 6D). Moreover, drought caused a significant increase in the expression of *Ferritin 1* in the roots relative to controls. Furthermore, FeEDTA supplemented with drought-stressed plants caused a significant increase in the expression of *Ferritin 1* in the roots compared to drought-exposed plants. Control plants treated with FeEDTA showed similar expression patterns of *Ferritin 1* in the roots to those of control plants (Fig. 6E).

**Fig. 6.**
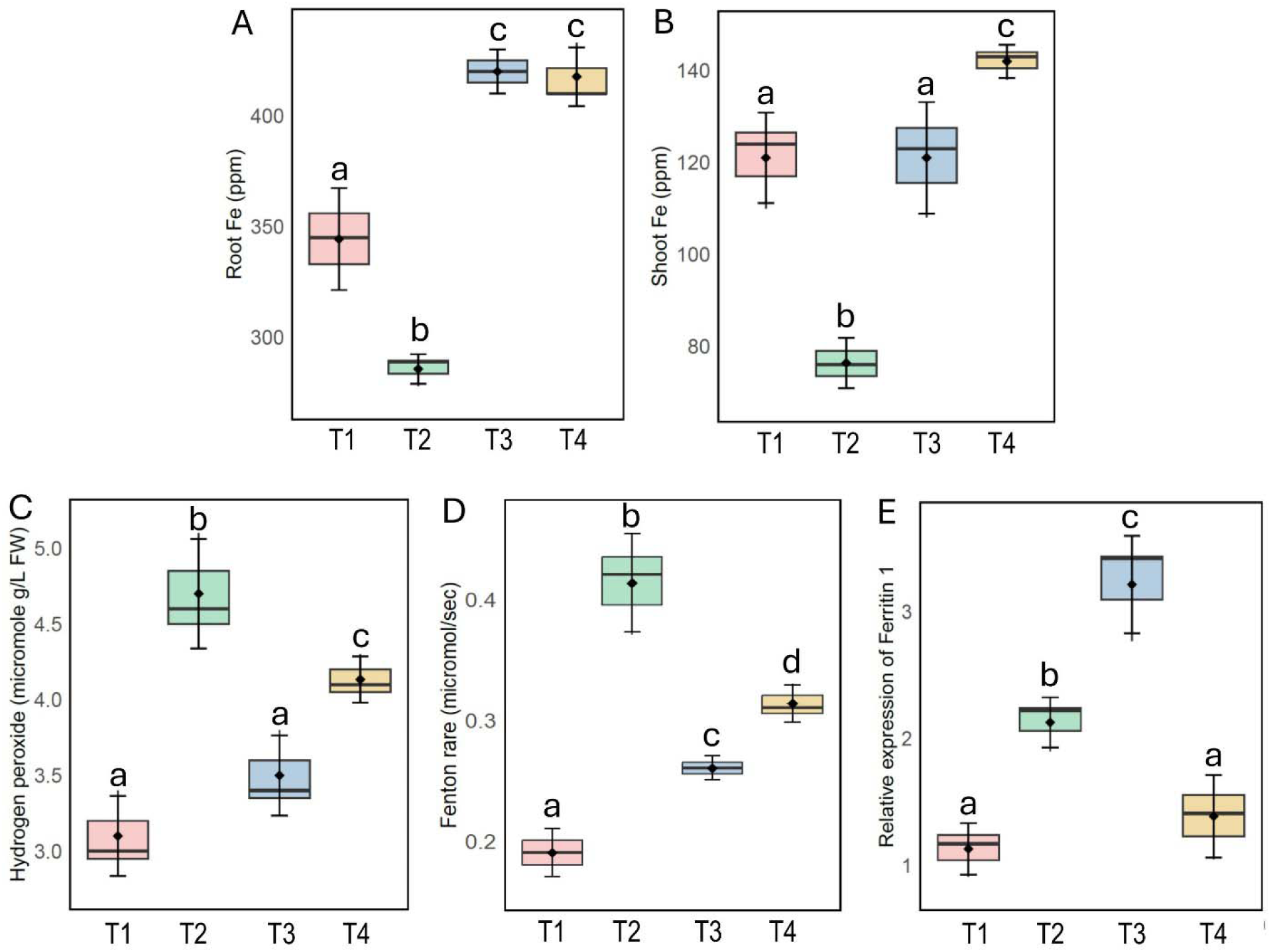
Changes in root Fe (A), shoot Fe (B), root hydrogen peroxide (C), root Fenton reaction rate (D) and relative expression of *Ferritin 1* in the roots (E) of sorghum cultivated with different drought and Fe treatments (T1: control, T2: drought, T3: drought + FeEDTA, T4: FeEDTA). The data presented are means with standard deviations (*n* = 3) of individual replicates). Statistically significant differences among the treatments at a *p* < 0.05 level are indicated by different alphabetical letters above the bars, where applicable. Plants were cultivated in a growth chamber for 4 weeks.

## Discussion

Drought stress can significantly impact sorghum growth and development. Understanding transcriptional and microbial shifts during the vegetative stage is crucial to conceptualizing strategies to enhance drought resilience in this vital crop. In this study, sorghum plants cultivated under drought conditions exhibited a significant improvement in plant biomass, leaf chlorophyll score, and relative water content when supplemented with exogenous Fe in the soil. Drought stress causes considerable damage to photosynthetic pigments, leading to chlorophyll degradation, which is considered a symptom of oxidative stress that causes growth retardation in plants (Ma et al., 2020; Rahdari et al., 2012). The reduced chlorophyll content under drought conditions may trigger the inactivation of photosynthesis and decreased transpiration efficiency in plants (Fahad et al., 2017). We also observed a decline in gas exchange parameters in sorghum exposed to drought. Of the gas-exchange parameters, stomatal conductance is most significantly affected by low water availability, because drought-stressed plants typically close their stomata to conserve water (Marchin et al., 2022; Farooq et al., 2009). While this response helps conserve water, it can also affect other physiological processes, such as photosynthesis and overall plant growth. In addition, drought conditions lead to a reduction in water use efficiency and transpiration rate, ultimately resulting in increased leaf and canopy temperatures (Turner et al., 2001). The morphological and physiological changes are coupled with the nutrient relations of plants exposed to drought stress (Ge et al., 2012). Furthermore, the composition and activity of soil microbial colonies are adversely impacted by soil water deficit, ultimately disrupting plant nutrient relations (Schimel et al., 2007). Previous studies showed that plants modify the length and surface area of their roots, as well as alter their architectural structure, to enhance the capture of less mobile nutrients (Lynch and Brown 2001). In this study, the decline of several nutrient elements in the roots of drought-exposed sorghum, particularly Fe and Mn may aggravate the reduction in bacterial richness as evident by the Observed richness in ASV. Previous studies have also shown that drought delays microbiome development and alters root metabolite profiles (Xu et al., 2018; Caddell et al., 2023).

The differential expression of genes associated with mineral homeostasis, hormone signaling, redox regulation, and osmotic adjustment in sorghum under drought stress reflects the plant’s multifaceted response to mitigate stress-induced damage and maintain growth. The upregulation of *Ferritin 1* (*Sobic.005G004500*), *Nitrate transporter 1* (*Sobic.009G119200*), and *Potassium transporter 12-related* (*Sobic.002G168800*) under drought conditions indicates that sorghum actively manages nutrient homeostasis to sustain metabolic activity and protect cellular structures. Ferritin, a key Fe-storage protein, reduces oxidative stress by sequestering excess Fe, thereby decreasing ROS formation (Ravet et al., 2009). Similarly, enhanced nitrate and potassium transporter expression suggests increased nutrient acquisition to counteract drought-induced nutrient limitations (Xu et al., 2021). We also found that elevated Fe resulted in a higher accumulation of Fe in the roots compared to the shoot. This was further substantiated by an increased expression of *Ferritin 1* in the root tissues when exogenous Fe was added to the drought-stressed plants, suggesting enhanced storage in shoot versus root under these conditions. Concurrently, H_2_O_2_, a common indicator of ROS damage under abiotic stress, substantially decreased in roots of drought-exposed sorghum with exogenous application of Fe in soil. Excess Fe increases ferritin abundance, enabling the storage of Fe, which contributes to the protection of plants against oxidative stress (Yadav et al. 2017; Reyt et al. 2015). Furthermore, the overexpression of the native ferritin gene *MusaFer1* enhances both Fe content and oxidative stress tolerance in transgenic banana plants (Parveen et al. 2016). The integration of Fe homeostasis with the generation of ROS is evident, and the distribution of Fe at the root tip plays a role in controlling root growth (Reyt et al. 2015).In another study, the regulation of Fe homeostasis through vacuolar Fe transporters and ferritin was found to enhance Fe accumulation in sorghum seeds that were from plants subjected to drought stress (Araki et al. 2022). This suggests that the increased Fe accumulation in the roots is associated with the induction of ferritin. This induction may play a role in reducing H_2_O_2_ levels and ensuring an adequate supply of Fe throughout the plant, which could restore chlorophyll synthesis and growth parameters in drought-exposed sorghum.

Furthermore, drought significantly altered hormone signaling, as evidenced by the upregulation of *Ethylene-insensitive protein 3* (*Sobic.004G181900*) and genes involved in gibberellin and auxin pathways, such as *Gibberellin 2-oxidase* (*Sobic.003G309800*) and *Auxin-responsive protein* (*Sobic.004G345600*). In addition, the modulation of auxin-related genes further highlights the importance of maintaining root architecture to optimize water and nutrient uptake (Sharma et al., 2020). Ethylene-insensitive proteins may also play a role in modulating stress responses by altering downstream transcription factors, as demonstrated in Arabidopsis under abiotic stress (Sakamoto et al., 2008). The interaction of auxin with ABA and ethylene often shifts the response toward stomatal closure as drought stress progresses (Lu et al., 2019). Consistently, our LC-MS analyses revealed the induction of ABA in the roots of sorghum supplemented under drought conditions. In the apoplast of *Arabidopsis thaliana*, ABA acts as a signal to induce stomatal closure by binding to receptors on guard cells, triggering a cascade of ion fluxes and turgor loss that minimize water loss during drought (Kuromori et al., in 2010 and 2014). Furthermore, *gibberellin 2-oxidase*, responsible for gibberellin degradation, may suppress growth-related processes to conserve resources, consistent with findings from drought studies on cereal crops (Shohat et al., 2021a). Drought-induced deactivation of gibberellins in guard cells promotes stomatal closure during the initial stages of drought in tomatoes (Shohat et al., 2021b). Interestingly, the levels of jasmonic acid significantly decreased in the roots of sorghum due to drought. Since plants prioritize drought survival strategies, such as maintaining water balance and minimizing cellular damage, the biosynthesis of jasmonic acid, which requires significant metabolic energy and carbon skeletons, might be downregulated to conserve resources in drought-exposed sorghum. Furthermore, drought stress strongly induces ABA in plants, as evidenced by our study. However, ABA often antagonizes JA signaling, potentially reducing jasmonic acid levels in plants, in response to biotic and abiotic stresses (Anderson et al., 2004). Our findings are consistent with the role of ABA as the primary drought-responsive hormone, though JA’s role in modulating root architecture and defense mechanisms under limited water availability warrants further exploration in sorghum.

Genes involved in redox regulation, including *Glutaredoxin* (*Sobic.003G259600*) and multiple *Glutathione S-transferases* (GSTs), were highly upregulated under drought stress. These genes play crucial roles in mitigating oxidative damage caused by ROS accumulation during water deficit (Chen et al., 2012). GSTs are known to detoxify harmful by-products of oxidative stress, such as lipid peroxidation products, and their induction is a hallmark of drought tolerance (Nahar et al., 2016). The expression of redox-regulating genes demonstrates the plant’s investment in maintaining cellular redox balance under drought. We also found the regulation of several genes related to water adjustment in the roots of sorghum exposed to drought. The upregulation of *Aquaporin* (*Sobic.009G075500*), *Dehydrin* (*Sobic.009G161700*), and *Glutamate decarboxylase 2* (*Sobic.007G152800*) suggests sorghum’s osmotic adjustment mechanisms to maintain cellular turgor. Aquaporins facilitate water transport across membranes, aiding in water retention during drought (Maurel et al., 2015). Dehydrins, known as late embryogenesis abundant (LEA) proteins, stabilize membranes and proteins under water-deficit conditions (Kosová et al., 2014). The induction of these genes highlights the importance of osmotic balance in maintaining physiological function during stress.

The downregulation of key genes in sorghum under drought stress may highlight the potential vulnerabilities in critical biological pathways but also point out the possible ways in which an unproductive root (i.e., one that is not taking up water or other nutrients) might be shut down or even sacrificed by the plant in order to concentrate nutrient discovery by the most productive roots. In the category of mineral uptake, the repression of *Zinc transporter* (*Sobic.007G075700*) and *Sulfate transporter ATPase* (*Sobic.006G124400*) indicates impaired nutrient acquisition, which may exacerbate stress-induced metabolic imbalances (Marschner, 2012). Similarly, the downregulation of hormone signaling genes, such as *Gibberellin 20-oxidase 2* (*Sobic.003G375900*) and *Auxin-responsive protein IAA6* (*Sobic.009G051600*), suggests a suppression of growth-related hormonal pathways, potentially leading to inhibited root and shoot development under water deficit conditions (Nelissen et al., 2018; Sharma et al., 2020), which may be vital in order to abandon these roots. Furthermore, the reduced expression of aquaporin transporter genes (*Sobic.003G105100* and *Sobic.004G251600*) likely compromises water transport and cellular turgor maintenance, further contributing to drought-mediated wilting and reduced physiological activity (Maurel et al., 2015). Additionally, the suppression of *Glutaredoxin* (*Sobic.003G259500*), a critical redox regulator, implies diminished capacity to mitigate oxidative stress caused by reactive oxygen species (ROS), exacerbating cellular damage (Noctor et al., 2012). Furthermore, the downregulation of genes associated with symbiotic signaling, such as *Phenylalanine ammonia-lyase* (*Sobic.004G220300*) and *Chalcone synthase* (*Sobic.006G103100*), may indicate a disruption in the plant’s ability to establish or maintain effective symbiotic relationships under stress conditions, potentially leading to reduced synthesis of key metabolites required for signaling pathways and mutualistic interaction dynamics in drought-exposed sorghum. Studies indicate reduced production of signaling compounds vital for microbe-plant interactions, potentially impairing nutrient acquisition through microbial symbiosis (Mitra et al., 2021; Mhlongo et al. 2018). These findings highlight how drought modulates key genes in symbiotic support, stress regulation, and nutrient relations, emphasizing the context-dependent roles of gene expression in response to drought in sorghum.

Drought stress causes a major compositional change in the rhizosphere and endophytic communities (Xu et al., 2021; Ma et al., 2020). In this study, amplicon sequencing data showed a reduction in both the diversity and richness of bacterial microbial communities, but not of fungi, in the roots of sorghum under drought conditions. In our study, fungal communities were dominated by *Talaromyces* under both experimental treatments. *Talaromyces* are primarily known for their role in the decomposition of organic matter (Heo et al., 2019). We found that *Talaromyces* were substantially enriched under drought, though these conclusions lack firm statistical support. The greenhouse conditions of this study were apparently selected for a narrower range of fungal taxa that had variable abundance responses to drought than seen for bacteria. Fungi are generally agreed to be more drought resistant than bacteria (Sun et al., 2020), and the stochasticity between replicates likely points to their being little selection pressure within this experiment from either the soil environment caused by drought, or from the plant in response to any beneficial associations with this narrow set of fungal taxa. In a recent study (Kaur and Saxena, 2023), *Talaromyces purpureogenus* has been identified as a drought-tolerant endophyte with significant plant growth-promoting attributes. When applied as a bioinoculant to wheat seedlings, *T. purpureogenus* significantly improved various physiological and biochemical growth parameters under both normal and drought-stressed conditions. Similarly, research by Pang et al. (2020) highlighted the drought tolerance of *Talaromyces purpureogenus*, which exhibited improved growth in rice plants under 10% polyethylene glycol (PEG)-induced drought stress (Pang et al., 2020). In our ITS analysis, we found a significant decrease in the relative abundance of *Malassezia* which is not commonly recognized as a typical endophytic fungus. The imposition of drought is a highly selective force on the fungal microbial community, possibly through encouraging the proliferation of a few putatively tolerant species at the expense of other members of the community or a reduction in those intolerant members leaving only tolerant ones surviving without associated abundance shifts. The nature of amplicon sequencing cannot readily unpick the difference between these scenarios. It should be noted, however, that severe drought is likely to kill some root tissues, and that this necrotic process might be enhanced by a plant looking to concentrate on its most productive roots (Gill and Jackson, 2000). Hence, fungi that remove unproductive roots as part of a general decomposition process might have more available root tissue to decompose, and thus become more prominent, under drought conditions.

When considering the bacterial community drought response, drought conditions appear to favor the proliferation of *Burkholderia-Caballeronia-Paraburkholderia* in the roots of sorghum. The taxonomy assignment “*Burkholderia-Caballeronia-Paraburkholderia*” reflects the complexities of microbial taxonomy, particularly within the *Burkholderia sensu lato* group, which has undergone significant reclassification in recent years (Bach et al., 2023). However, the genera *Burkholderia*, *Caballeronia*, and *Paraburkholderia* encompass a diverse group of bacteria known for their plant growth-promoting properties, including enhancing drought tolerance (Ahmad et al., 2022). *Paraburkholderia phytofirmans* strain PsJN produces phytohormones such as indole-3-acetic acid (IAA), which stimulate root elongation and branching, thereby improving water uptake under drought conditions (Donoso et al., 2016). Furthermore, auxin is found to be a key determinant in mycorrhizal symbiotic association with plants (Kabir and Bennetzen, 2024; Ludwig-Müller and Güther, 2007). The genera belonging to the Burkholderiaceae family are known for containing a diverse set of taxa capable of plant symbiosis and functions in plant growth, stress resilience, and nitrogen fixation (Pal et al., 2022; Esmaeel et al., 2020). *Burkholderia* species have been well documented to be associated with an increase in the drought stress resilience of host plants in inoculation studies (Naveed et al., 2014; Naveed et al., 2014; Huang et al., 2017; Tallapragada et al., 2016). Studies have also reported that switchgrass inoculated with *Burkholderia* were more photosynthetically active, allocated more biomass, and produced more tillers than uninoculated plants (Wang et al., 2015; Kim et al., 2012). *Burkholderia* may alleviate drought stress by producing Fe-chelating compounds (siderophores) that help plants survive in environments with low Fe levels (Pal et al., 2022; Barrera-Galicia et al., 2021). Both *Paraburkholderia* and *Burkholderia* have also been identified as inorganic phosphate-solubilizing bacteria (Damo et al., 2022). In light of this evidence, our findings underscore the potential application of *Burkholderia-Caballeronia-Paraburkholderia* species as bioinoculants to enhance drought tolerance in sorghum, offering a sustainable solution to mitigate the adverse impacts of water deficit in soil.

The decline in *Rhizobium*, *Herbaspirillum*, and *Sphingomonas* that we observed under drought conditions is likely due to reduced soil moisture, altered root exudation patterns, and/or oxidative stress, which collectively create a less hospitable environment for these potentially beneficial microbes (Jones et al., 2019; Ricks et al., 2023). Although sorghum roots do not host *Rhizobium* for nitrogen fixation, sorghum may interact with other beneficial microbes in the rhizosphere, such as diazotrophic bacteria that can fix nitrogen and contribute to plant growth. Furthermore, drought-driven decreased accumulation of soil iron by sorghum may disrupt bacterial metabolic processes. These stresses may alter sorghum root exudation patterns, making them less favorable for sustaining beneficial microbes, as we observed in drought-exposed sorghum. *Herbaspirillum* and *Sphingomonas*, recognized for their roles in phytohormone production and nutrient solubilization (Grillo-Puertas et al., 2021; Yang et al., 2014), also exhibited reduced population density in response to drought, thereby potentially impairing their plant growth-promoting functions. Metze et al. (2023) showed that drought inhibited the growth of over 90% of bacterial taxa, but favored the growth of specialized taxa, including Actinomycetota such as *Streptomyce*s. In this study, we observed a less significant increase in *Streptomyces* abundance in response to drought stress than seen in previous studies conducted in soil systems (Metze et al., 2023; Xu et al., 2021; Fitzpatrick et al., 2018). As our experiment was conducted in greenhouse pots, and not in the field as per the other referenced studies, some differences in outcomes would be expected because our observed initial community is likely a subset of the field community (Kushwaha et al., 2024). For example, Forero et al. (2019) showed that different plant-soil feedback is observed between greenhouse and field trials even when greenhouse pots are inoculated with field soil. Our findings suggest that the drought stress we imposed had a minimal impact on the overall functional composition of bacterial and fungal communities, compared to the major physiological and transcriptional changes observed. This aligns with studies that propose microbial functional stability as a mechanism for ecosystem resilience to environmental stress (Evans & Wallenstein, 2014; Griffiths and Philippot, 2013). However, the divergent capacities of bacterial communities to respond functionally under drought were evident in the predictive accuracy of the random forest models. Bacterial functions were effectively predicted (error rate of 12.5%), indicating that bacterial communities may exhibit a more defined or deterministic functional response to drought in sorghum.

While bacterial communities exhibited clear functional shifts under drought conditions, the fungal communities showed minimal changes in diversity and composition, with their functional responses being less distinct. The limited functional predictability for fungal communities, as evidenced by the high error rate in random forest models, may be attributed to their lower diversity under drought conditions and potential functional redundancy among dominant taxa, which could mask distinct functional shifts in response to drought stress. Identifying functional groups associated with bacterial communities highlights their distinct responses under drought and control conditions, showcasing microbial adaptation strategies. Functional groups such as chorismate metabolism, glycolysis, and protochatechuate degradation I was more active in the control conditions, reflecting bacterial contributions to primary metabolism and carbon cycling under stable conditions. Conversely, functional groups like clavulanate biosynthesis, phototrophy, and photosynthetic cyanobacteria were associated with drought, highlighting shifts in bacterial functional strategies to adapt to water scarcity. These results are consistent with findings by Lennon et al. (2012), which suggest that bacterial communities can reconfigure their functional potential to maintain ecosystem processes under drought stress. The integration of the MetaCyc and FAPROTAX databases provided complementary insights into these functional dynamics. While MetaCyc predicted most groups, FAPROTAX identified additional drought-associated functions such as methyltrophy and nitrate reduction. These functions are critical for nutrient cycling and drought resilience, as methylotrophic activity and nitrate reduction can mitigate resource limitations (Torres Vera et al., 2024; Deng et al., 2024). These results highlight the importance of combining multiple annotation databases to obtain a comprehensive understanding of microbial functional shifts under environmental stress. Overall, the results suggest that bacterial communities exhibit greater functional plasticity in response to drought stress compared to fungal communities. This finding emphasizes the need for future studies to explore the underlying mechanisms driving these functional responses, such as shifts in microbial taxa, substrate availability, and resource competition.

## Conclusion

Taken together, this study underscores the complex responses of sorghum exposed to drought stress at both the transcriptional and microbial levels. The upregulation of genes related to nutrient homeostasis, hormone signaling, and redox balance demonstrates the plant’s ability to activate pathways essential for maintaining physiological processes under stress. Drought-induced genes such as *Ferritin 1*, *Fe dehydrogenase* and *Nitrate transporter 1* play critical roles in mitigating oxidative damage and ensuring nutrient uptake, while aquaporins and dehydrins highlight the importance of osmotic regulation in sustaining cellular function during water deficit (Fig. 7). Also, drought tolerance sorghum may be associated with the induction of *Ferritin 1*, which resulted in a reduction in Fenton rate and H_2_O_2_. These responses reflect the adaptive strategies of sorghum to survive under drought conditions. In parallel, drought significantly shaped the root-associated microbiome, favoring specific bacterial taxa like *Burkholderia-Caballeronia*-*Paraburkholderia*, known for their contributions to plant growth through auxin production, phosphate solubilization, and Fe chelation. The enrichment of functional groups such as phototrophy and nitrate reduction further illustrates the role of microbiome dynamics in supporting nutrient availability and plant resilience. These findings emphasize the potential of harnessing plant genetics and beneficial microbial consortia as key breeding targets to enhance sorghum’s drought tolerance. Future efforts should emphasize a breeding approach, prioritizing field-based validation and the integration of these insights into programs for developing elite sorghum genotypes and effective bioinoculants to strengthen sustainable agriculture in water-limited environments.

**Fig. 7.**
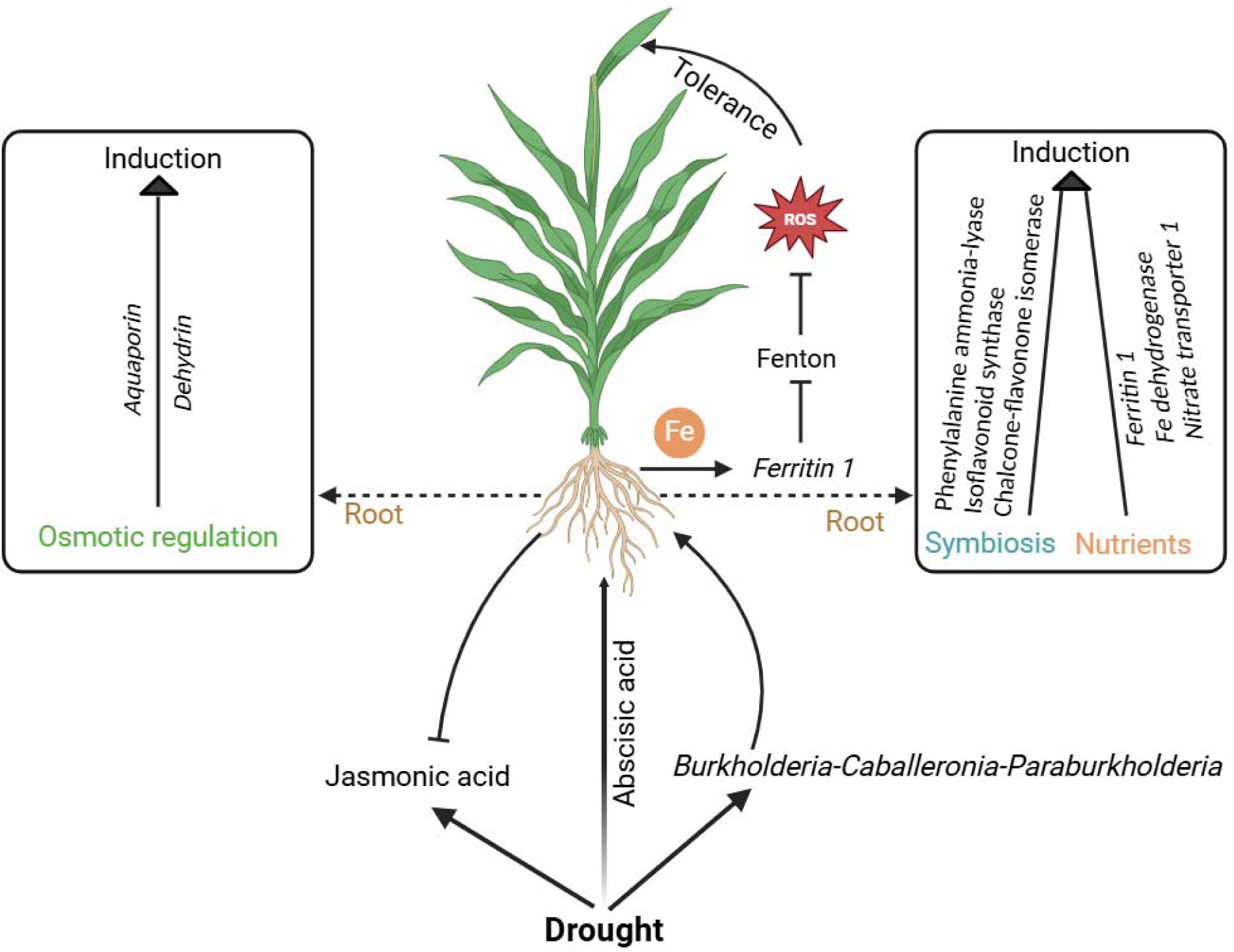
Shift in hormonal regulation, transcriptome and microbiome dynamics in sorghum exposed to drought. Drought stress enhances abscisic acid (ABA) levels, leading to osmotic regulation through the induction of *Aquaporin* and *Dehydrin*. Simultaneously, jasmonic acid (JA) levels are suppressed, possibly due to its antagonistic interaction with ABA. Furthermore, Fe supplementation drives *Ferritin 1* induction thereby reducing Fenton reaction-mediated reactive oxygen species (ROS) to induce drought tolerance. In response to drought, symbiotic and nutrient-related processes are upregulated, as evidenced by the induction of genes such a *Phenylalanine ammonia-lyase*, *Isoflavonoid synthase*, and *Chalcone-flavonone isomerase* (symbiosis-related), along with *Ferritin 1*, *Fe dehydrogenase*, and *Nitrate transporter 1* (nutrient acquisition-related). The root microbiome, including species such as members of th *Burkholderia-Caballeronia-Paraburkholderia*, potentially contributes to these adaptive responses by supporting nutrient uptake and stress resilience.

## Data availability

Illumina sequencing data was submitted to NCBI under the following accession numbers: PRJNA1128172 (RNA-seq), PRJNA1128337 (16S) and PRJNA1128338 (ITS).

## Acknowledgments

We express our gratitude to the Georgia Genomics and Bioinformatics Core (GGBC) and the Mass Spectrometry and Metabolomics Core of Michigan State University. This research received support from the US Department of Energy (grant DE-SC0021386), Georgia Research Alliance, and a startup grant (5SFAES-293007) from the University of Louisiana at Monroe.

## Author contributions

AHK conceived the idea for the study, conducted physiological assays, RNA-seq analysis, split-root assay, and prepared the draft manuscript. PBJ prepared the DNA library, analyzed the amplicon sequencing data, and wrote the microbial part of the results. MA crosschecked the RNA-seq data, performed gene enrichment analysis, and reviewed the discussion. JL carried out the bioinformatics analysis of microbial networks and functional groups. BA interpreted the microbial functional analysis section and critically reviewed the manuscript. SPLT provided significant contributions to the manuscript. JLB supervised the work, provided intellectual input, and critically revised the manuscript.

## Notes

### Competing Interest Statement

The authors have declared no competing interest.

### Summary of Updates

We have changed the title of the manuscript to be more emphasized on the main findings of the study.

